# MicrographCleaner: a python package for cryo-EM micrograph cleaning using deep learning

**DOI:** 10.1101/677542

**Authors:** Ruben Sanchez-Garcia, Joan Segura, David Maluenda, C.O.S. Sorzano, J.M. Carazo

## Abstract

Cryo-EM Single Particle Analysis workflows require from tens of thousands of high-quality particle projections to unveil the three-dimensional structure of macromolecules. Conventional methods for automatic particle picking tend to suffer from high false-positive rates, hurdling the reconstruction process. One common cause of this problem is the presence of carbon and different types of high-contrast contaminations. In order to overcome this limitation, we have developed MicrographCleaner, a deep learning package designed to discriminate which regions of micrographs are suitable for particle picking and which are not in an automatic fashion. MicrographCleaner implements a U-net-like deep learning model trained on a manually curated dataset compiled from over five hundred micrographs. The benchmarking, carried out on about one hundred independent micrographs, shows that MicrographCleaner is a very efficient approach for micrograph preprocessing. MicrographCleaner (micrograph_cleaner_em) package is available at PyPI and Anaconda Cloud and also as a Scipion/Xmipp protocol. Source code is available at https://github.com/rsanchezgarc/micrograph_cleaner_em.

## 1. Introduction

Cryogenic-Electron Microscopy (cryo-EM) Single Particle Analysis (SPA) has recently become a powerful technique for the determination of macromolecular structures achieving, in many cases, atomic resolutions. SPA consists of a set of complex and variable operations that, departing from thousands of particle projections, leads to the synthesis of electronic density maps of macromolecules. The massive number of particles that are needed for SPA has made of automatic particle picking one of the most influential steps in virtually all reconstruction workflows. Nevertheless, some problems intrinsic to the cryo-EM pipelines, such as low signal-to-noise ratio and the presence of high contrast artifacts and contaminants in the micrographs, degrades the performance of particle picking algorithms (Vargas et al., 2013; Zhu et al., 2004) and leads to the addition of false positive particles in SPA workflows. This problem can be mitigated trough different algorithms that clean and remove incorrectly selected particles after automatic picking (Sanchez-Garcia et al., 2018; Vargas et al., 2013).

One of the most common shortcomings observed during automatic picking is the attraction of these methods to select grid carbon spots, especially at the hole edges. Due to its relevance, some algorithms have been designed to prevent particle selection in those regions. For example, the em_hole_finder program, included in the Appion package (Lander et al., 2009) is based on morphological image processing operations to compute masks around carbon holes. Similarly, EMHP (Berndsen et al., 2017) was designed to perform a similar task through image filtering and thresholding operations followed by a circle fitting procedure. Although very useful when grid edges are clearly visible, both approaches struggle in those cases where high contrast contaminations are present in micrographs. Moreover, both of them require human supervision to determine the presence of carbon in the micrographs and to set some user-defined parameters. As a result, its applicability is limited to supervised scenarios.

More recently, deep learning particle pickers have been developed with the aim of improving picking accuracy. (Bepler et al., 2019; Wagner et al., 2019; Wang et al., 2016; Zhang et al., 2019; Zhu et al., 2017). These new particle pickers are more robust to false positives and most of them have been explicitly or implicitly designed to avoid carbon areas and large contaminants. One of such explicitly designed particle pickers is included in the Warp package (Tegunov and Cramer, 2019). Thus, the Warp picking algorithm approaches the problem of particle picking performing a pixel-wise classification -segmentation-of the micrographs in which one of the possible categories is undesirable region.

Nevertheless, despite these developments, conventional particle pickers are still the preferred choice in recent publications (Gilman et al., 2019; Hiraizumi et al., 2019; Jain et al., 2019; Molina et al., 2019; Stone et al., 2019; Yan et al., 2019). Although it is likely that deep learning particle pickers will become increasingly popular, they are not perfect and different situations will require different approaches, so conventional particle pickers, especially those based on templates, will probably remain popular.

Following this line, and with the aim of improving classical particle pickers and complementing deep-learning-based ones, we have developed MicrographCleaner, a fully automatic, easy-to-install and easy-to-use deep learning solution that performs a pixel-wise classification of micrographs into two categories, desirable and undesirable regions for picking. Like Warp particle picker, MicrographCleaner relies on one of the most extended network architectures (Ronneberger et al., 2015), the U-net, but the different choices in important parameters result, in turn, in quite different levels of performance. Thus, according to our benchmarking, MicrographCleaner is not only able to provide a more robust and accurate solution for carbon detection than earlier methods, but it is also able to improve the detection of other types of contaminants, such as ice crystals or ethane bubbles.

## 2. Material and methods

### 2.1 Algorithm

MicrographCleaner computes binary segmentation of micrographs with the aim of delineating optimal regions for particle picking and isolating those areas containing high-contrast contaminants and other artifacts. To that end, MicrographCleaner implements a U-net-like architecture (Ronneberger et al., 2015). Our model, carefully selected after a cross-validation process, consists of 5 downsampling blocks followed by 5 upsampling blocks with 32, 64, 128, 256, and 512 kernels per block respectively. Further details are described in Supplementary Material S1 and S4.

Neural network training was carried out during 200 epochs using the Adam optimizer and a combination of perceptual loss (Johnson et al., 2016) and weighted binary cross-entropy (Falk et al., 2019). Data augmentation was performed during training. See Supplementary Material S2 and S4 for more details.

### 2.2. Dataset and preprocessing

MicrographCleaner was trained on a dataset of 539 manually segmented micrographs collected from 16 different EMPIAR (Iudin et al., 2016) entries. The evaluation was performed on an independent set of 97 micrographs compiled from two EMPIAR projects and another two in-home projects (see Supplementary Material S5). Both training and testing sets micrographs include examples of clean, carbon-containing, contamination-containing and aggregation-containing areas as well as mixed ones that were labeled by an expert.

Before micrographs are fed to the network, a previous normalization step is required to adjust the different intensity scales and sizes of micrographs. Thus, all micrographs are normalized in both intensity and size using a robust scaling strategy and a constant particle size donwsampling (see Supplementary Material S3). Finally, due to GPU memory limitations, the full downsampled micrograph is processed in chunks using a sliding window approach of overlapping patches of size 256×256.

### 2.3. Evaluation metrics

As evaluation criterium, we have computed the Intersection over Union (IoU) metric between the network predictions and the manually curated masks and averaged it for all micrographs in the testing set (mIoU). Consequently, mIoU is defined as:

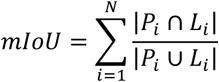

where i is the testing micrograph index, N the number of testing micrographs and Pi and Li are, respectively, the predicted mask and the manually curated mask for testing micrograph i.

### 2.4 Package

MicrographCleaner has been implemented as an easy-to-install and easy-to-employ Python 3.x package. Thus, the command line tool can be automatically installed from Anaconda Cloud and PyPI repositories whereas the GUI version can be installed through the Scipion (de la Rosa-Trevín et al., 2016) plugin manager. The neural network was implemented using Keras (Chollet, 2015) package and Tensorflow (Abadi et al., 2016) backend. Micrograph preprocessing is carried out using the scikit-image (van der Walt et al., 2014) package.

## 3. Results

### 3.1 Carbon detection

In order to estimate carbon detection capability, we have taken the subset of the testing set micrographs in which all contain some carbon and we have executed MicrographCleaner on them, achieving a mIoU of 0.78833, which indicates a great agreement between the carbon areas manually curated and the predicted ones.

Additionally, we have compared MicrographCleaner with several carbon finder programs: em_hole_finder, EMHP and the Warp particle picker (WPP) (see Supplementary Material S6 for more details). Before entering into these comparisons, it is important to highlight that MicrographCleaner and the WPP, contrary to the others, are fast (in the order of seconds), parameter-free and they do not require manual intervention in order to determine whether or not carbon is present in a micrograph. Consequently, they can be employed in automatic pipelines and, thus, they are suitable for automatic Cryo-EM analysis at facilities. As it can be appreciated in Table 1 and in Supplementary Material SM1, deep learning-based methods are very well suited for this problem as both Warp and MicrographCleaner stand out from the others. Still, MicrographCleaner achieves the best performance of all them by a wide margin, improving results over the second best, WPP, by more than 20% in terms of agreement between masks predictions and ground truth.

**Table 1.** MicrographCleaner performance for carbon detection compared to other methods.

| Algorithm | mIoU | stdIoU |
| --- | --- | --- |
| MicrographCleaner | 0.78833 | 0.22939 |
| EMHP | 0.19805 | 0.21147 |
| `em_hole_finder` | 0.05691 | 0.04691 |
| Warp Particle Picker | 0.57297 | 0.23095 |

Notes: mIoU: mean Intersection over Union (mean fraction of agreement between predictions and ground truth); stdIoU: standard deviation Intersection over Union.

### 3.2. Undesirable regions and contaminants detection

MicrographCleaner evaluation for undesirable regions and small contaminants detection was performed comparing the predicted masks with the ground truth for all the testing micrographs. Under this test, MicrographCleaner achieved a mIoU value of 0.544. This score, although worse that the score for carbon detection, implies a good agreement between ground truth and predicted masks, especially when taking into account that the testing set contains clean micrographs examples together with carbon-containing and contaminated micrographs. Figure 1 shows the predictions for four different micrographs, illustrating that MicrographCleaner is capable of successfully detecting both contaminants and carbon.

**Fig 1.**
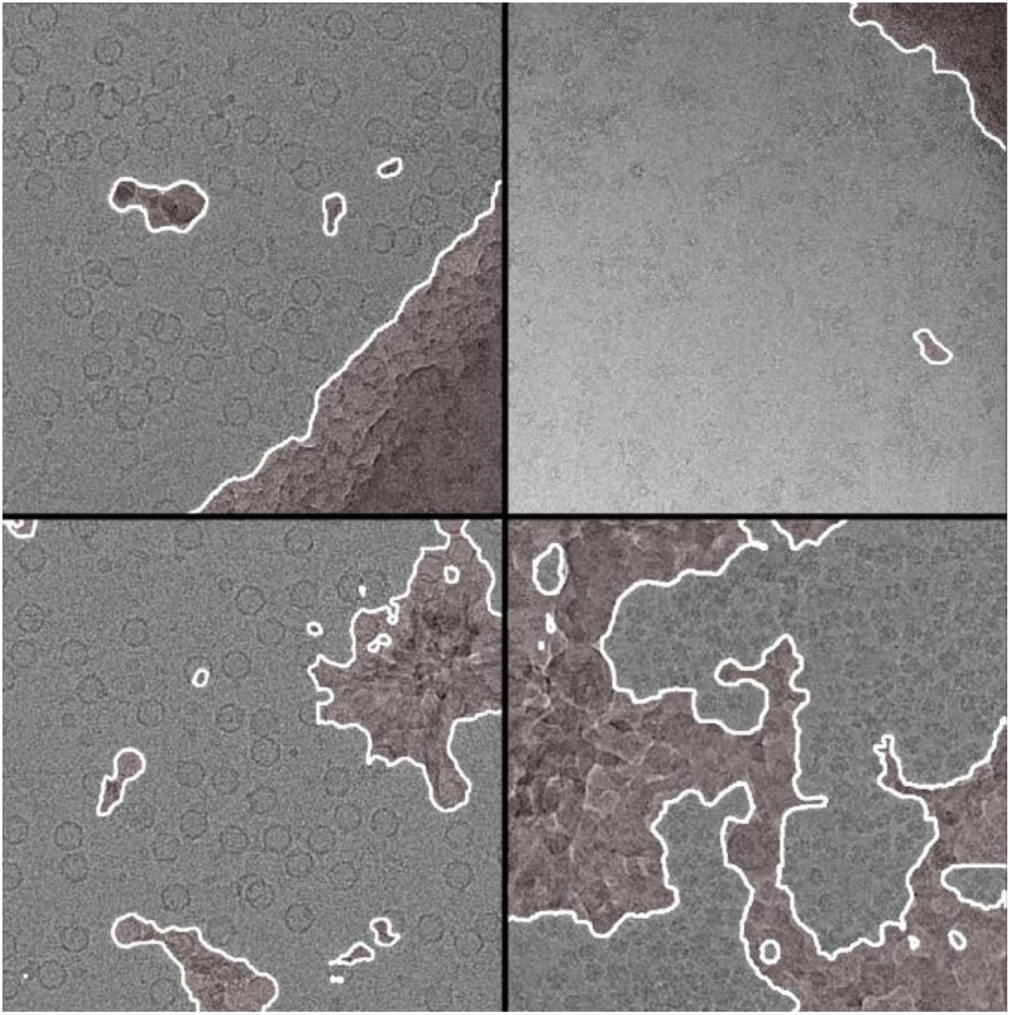
MicrographCleaner identifies non-suitable regions. Red shadowed regions correspond to micrograph areas labeled as “non-suitable” with 50% or more confidence. Top images show MicrographCleaner capability to detect carbon in the presence of contaminants. Bottom images show MicrographCleaner capability to detect a wide variety of different contaminants.

Additionally, we have also evaluated the global performance of WPP on the whole testing set, showing a mIoU of 0.331 and performing worse than MicrographCleaner for 77% of the micrographs. This supposes that the 20% better performance of MicrographCleaner over WPP for carbon detection is also maintained when contaminants detection is also considered. The predictions for some micrographs using both MicrographCleaner and WPP are shown in Supplementary Material Figure SM2.

### 3.3. Use cases

In this section we present two examples, not included in the training and testing sets, in which both traditional particle pickers and deep-learning-based pickers struggle discerning problematic regions and contaminants from clean regions and thus, they both could benefit from MicrographCleaner. As deep learning representatives, we have chosen Topaz (Bepler et al., 2019) and the Cryolo (Wagner et al., 2019) particle pickers. Both Cryolo and Topaz algorithms were trained using ten manually curated micrographs. Additionally, the Cryolo general model, that does not require any training, was also employed. Relion autopicker (Scheres, 2015) was chosen as the representative of traditional particle pickers. Further details can be found in Supplementary Material S8.

#### 3.3.1. EMPIAR-10156

The main difficulties for particle pickers that EMPIAR-10156 dataset (von Loeffelholz et al., 2018) presents is that it contains large areas of carbon (even more than 50% of the micrograph) and that the intensity of these areas is not uniform neither within an individual micrograph nor at the whole dataset. Thus, as it is illustrated in Figure 2, both the Relion and the Cryolo particle pickers (using a general model and a trained one) tend to pick particles located at the carbon region, whereas Topaz particle picker is able to avoid most of the carbon region but still selects many false positives at the edge.

**Fig 2.**
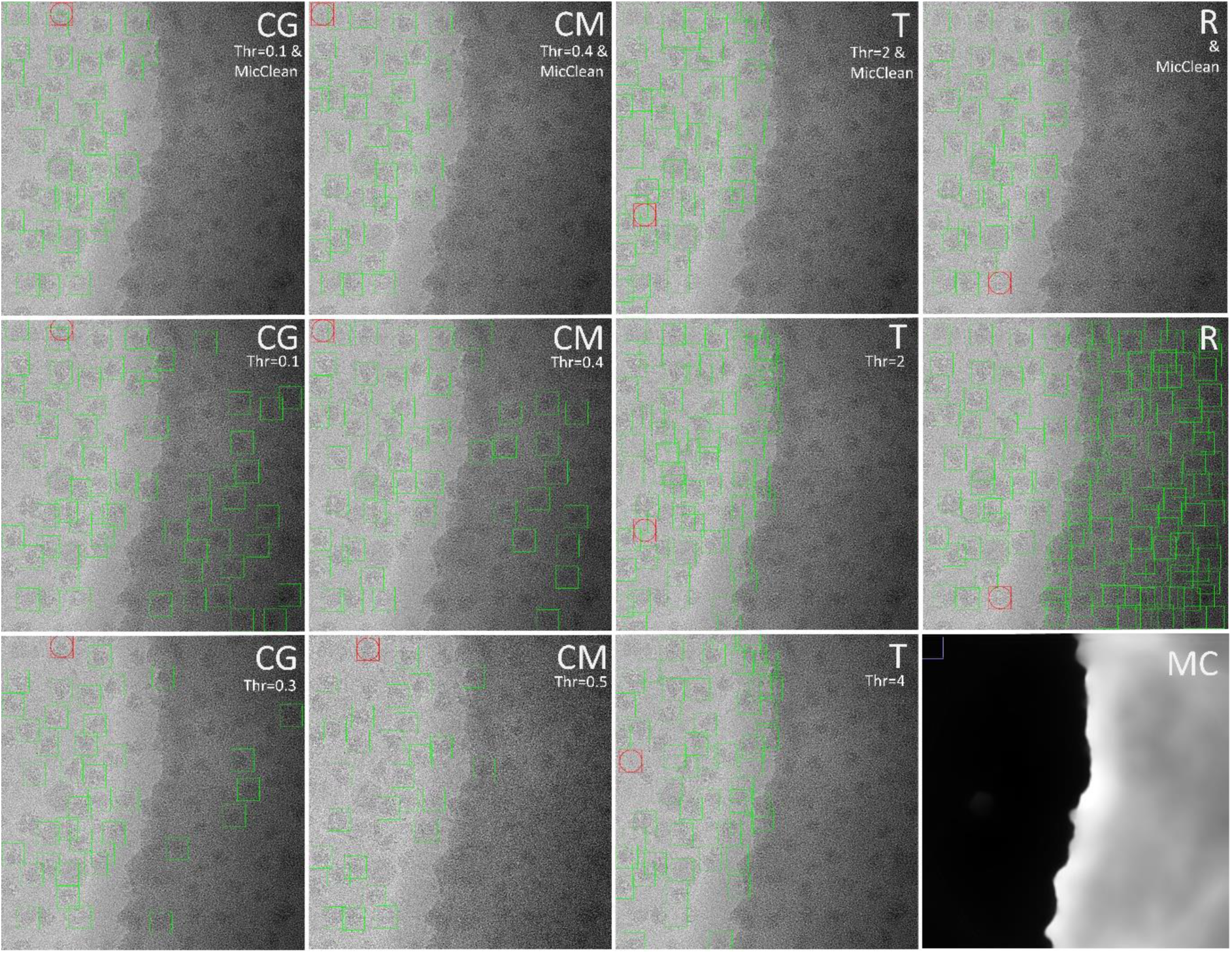
MicrographCleaner improves particle picking on EMPIAR-10156 dataset. Coordinates selected with Cryolo pretrained general model (CG), Cryolo manually trained model (CM), Topaz (T) and Relion autopicker (R) are respectively displayed in columns one to four. Top row images correspond to the remaining particles after applying MicrographCleaner mask (MC) to the low threshold Topaz, Cryolo general and Cryolo manual solutions and the Relion autopicker outcome. As it can be appreciated, MicrographCleaner removes the particles selected in the carbon area and its edge while preserving much more true positive particles than using stricter thresholds. Red box represents the lowest confidence particle according the picking algorithm.

It is interesting to consider that, although the number of particles picked at the carbon area/edge can be easily decreased using stricter thresholds, it comes at a cost of ruling out true positive particles. Thus, as it is shown in Figure 2, large enough thresholds for discarding most of the false positive particles cause the rejection of some true positive particles, which in the end translate to the precision/recall tradeoff in which most people favor the latter option aiming to remove false positives in successive steps. On the other hand, MicrographCleaner is able to mask out those false positive particles while not affecting the true positive ones, hence it can be used as a complement for any particle picker independently of threshold decisions. This behavior is illustrated in Figure 2, in which MicrographCleaner proposed solutions are better than the solutions obtained directly by the other methods at different thresholds. For more details see Supplementary Material S8.

#### 3.3.2. *EMPIAR-*10265

The EMPIAR-10265 dataset (Lee et al., 2019) is extremely challenging. In this dataset, the particles of most micrographs are difficult to visualize, whereas in some others they are easily recognizable (see Figure 3 and 4 respectively). Due to this profound disparity, the performance of the employed deep-learning-based methods is worse than in other datasets, and although they are able to avoid large contaminated regions, they still incorrectly select many small contaminants as particles, as it is illustrated in Figure 3 and 4. Again, like in the previous example, the number of selected contaminants can be reduced by increasing the threshold but, as a result, the total number of particles is severely reduced.

**Fig 3.**
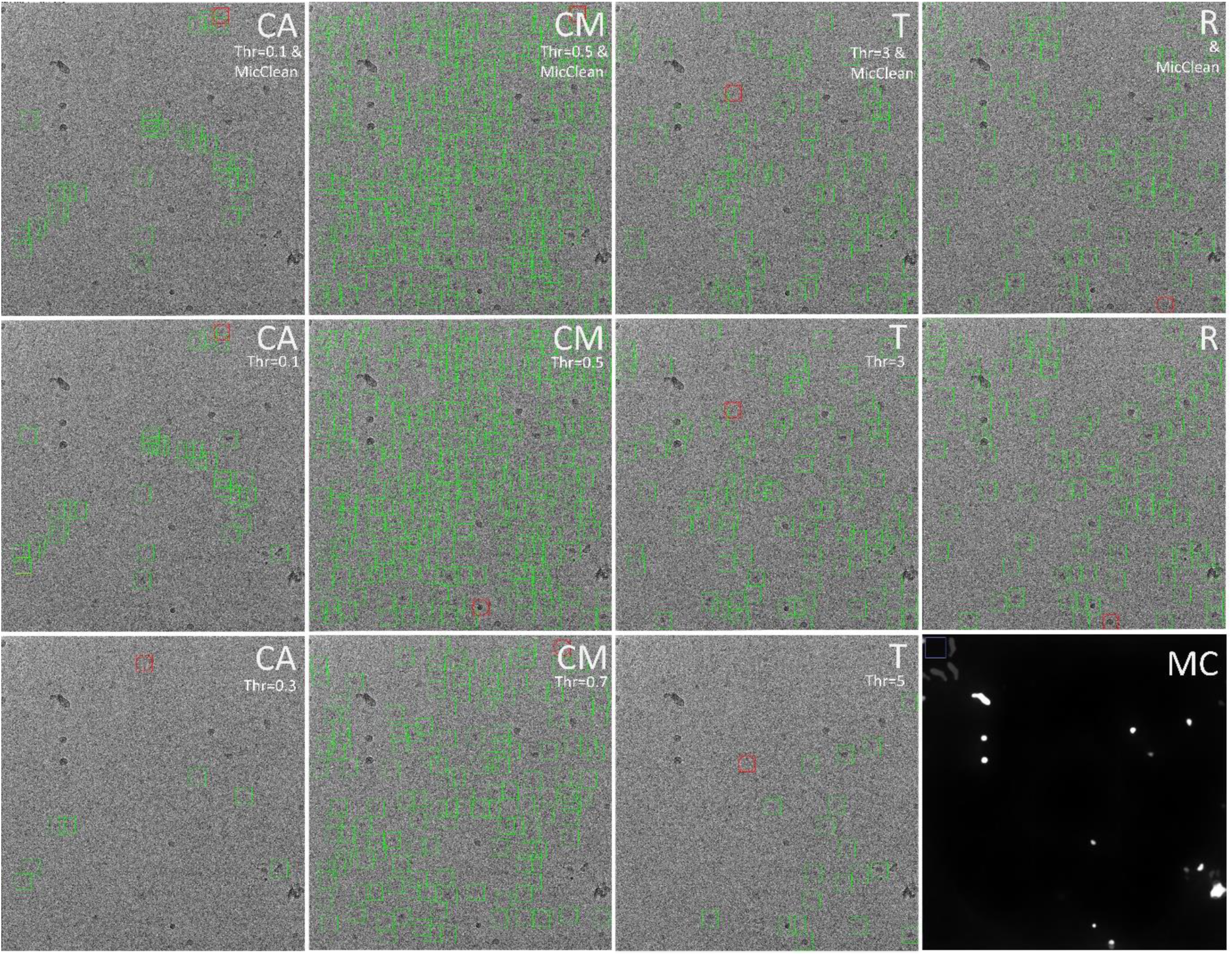
MicrographCleaner improves particle picking on EMPIAR-10265 dataset. Coordinates selected with Cryolo pretrained general model (CG), Cryolo manually trained model (CM), Topaz (T) and Relion autopicker (R) are respectively displayed in columns one to four. Top row images correspond to the remaining particles after applying MicrographCleaner mask (MC) to the low threshold Topaz, Cryolo general and Cryolo manual solutions and the Relion autopicker outcome. As it can be appreciated, MicrographCleaner removes many of the contaminants incorrectly selected as particles while preserving much more true positive particles than using stricter thresholds. Red box represents the lowest confidence particle according the picking algorithm.

**Fig 4.**
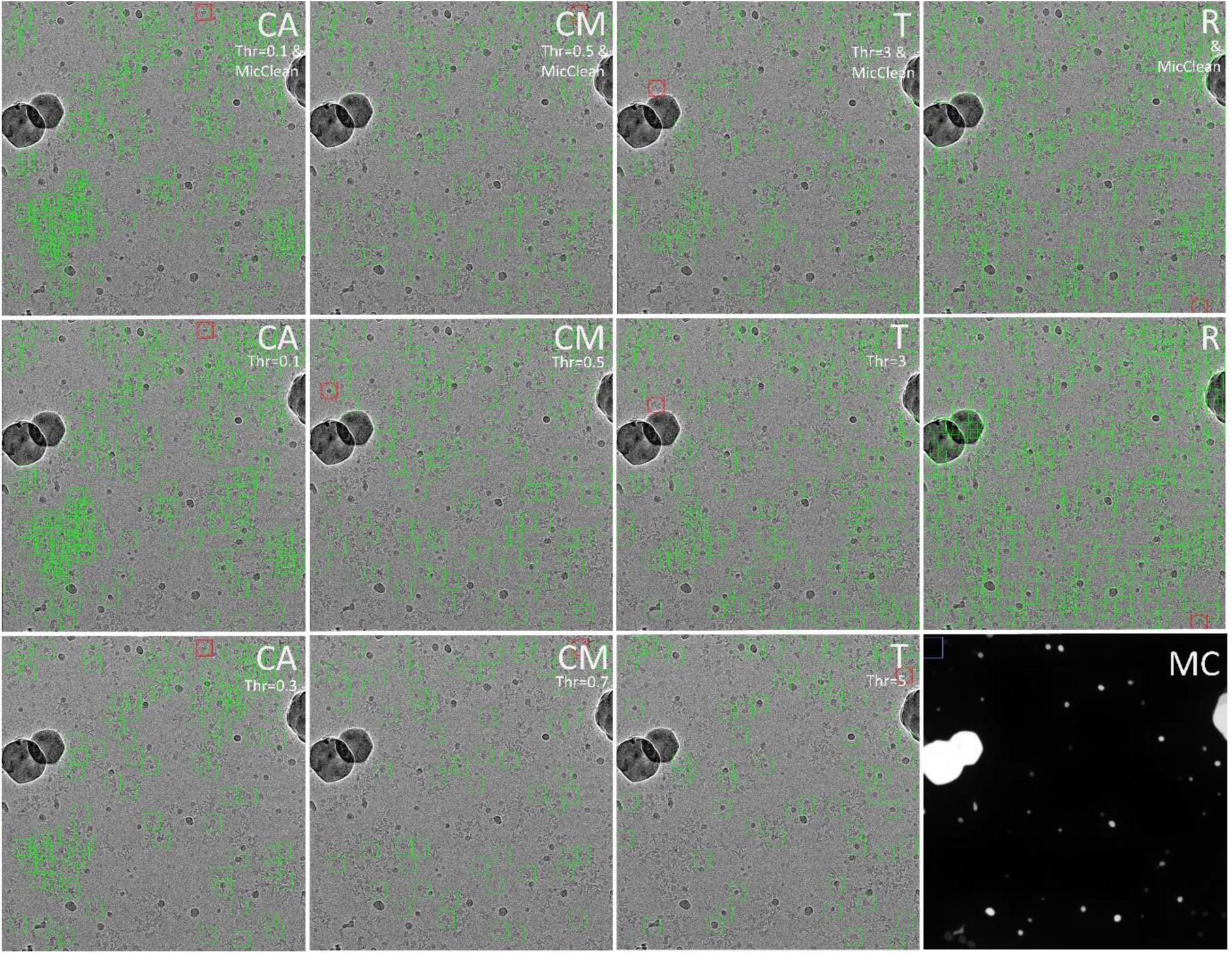
MicrographCleaner improves particle picking on EMPIAR-10265 dataset. Coordinates selected with Cryolo pretrained general model (CG), Cryolo manually trained model (CM), Topaz (T) and Relion autopicker (R) are respectively displayed in columns one to four. Top row images correspond to the remaining particles after applying MicrographCleaner mask (MC) to the low threshold Topaz, Cryolo general and Cryolo manual solutions as well as the Relion autopicker outcome. As it can be appreciated, MicrographCleaner removes many of the contaminants incorrectly selected as particles while preserving much more true positive particles than using stricter thresholds. Red box represents the lowest confidence particle according to the respective picking algorithm.

Thus, the process of threshold selection for this dataset is not trivial, as micrographs differ severely and thresholds that detect most of the particles in some micrographs discard many particles in others. As a consequence, manual inspection for each micrograph should be performed to obtain the best balance between the number of removed contaminants and total number of recovered particles. Alternatively, although still expensive, more micrographs could be manually picked in order to train further some of the methods.

On the other hand, when MicrographCleaner is applied to the particles that have been selected using a conservative threshold, more true positive particles can be recovered while ruling out most of the small contaminants that were incorrectly selected (see Figure 3 and 4). This ultimately improves the quality of the set of picked particles and also simplifies threshold selection, that can be set to more conservative values with the confidence that contaminants will be equally removed. See Supplementary Material S8 for additional information.

### 3.4. MicrographCleaner complements 2D-classification

Although the previous section demonstrates that MicrographCleaner is able to reduce false positive levels for many particle pickers, it could also be argued that this reduction is not of enormous impact as such a reduction will be equally achieved by the subsequent steps of the image processing workflow, especially, at the 2D-classification step. With the aim of testing this hypothesis, we have conducted one 2D classification analysis for each of the particle sets picked by the four particle pickers considered in Section 3.3.1 and we have compared the outcome of all of them with the particle sets processed with MicrographCleaner. Figure 5 illustrates the experiment for one of the picked sets of particles (see Supplementary Material S9 for additional information and other examples).

**Fig 5.**
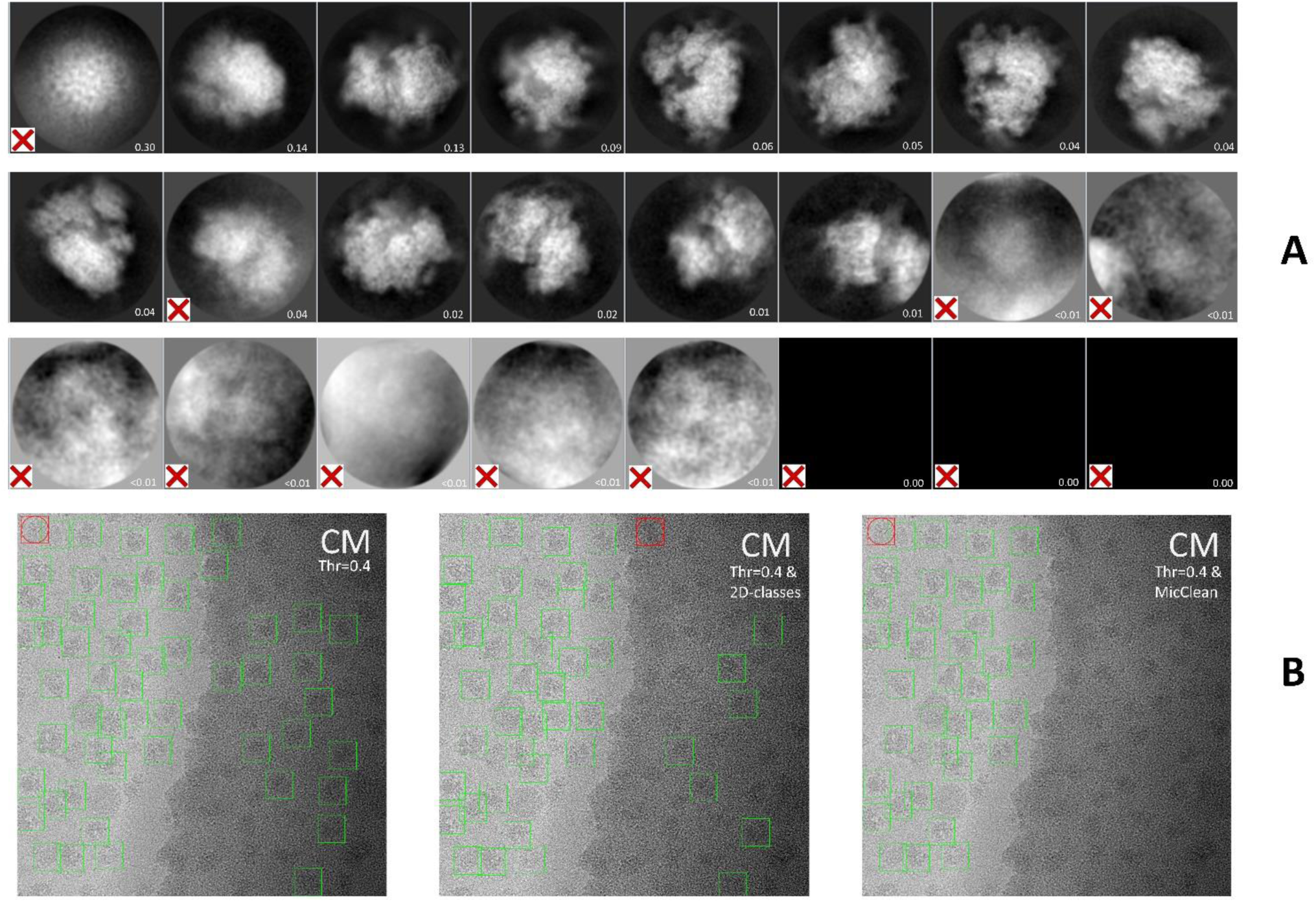
MicrographCleaner complements 2D-classification. A: Gallery of 2D averages obtained from the set of particles collected by Cryolo manually trained on EMPIAR-10265 dataset. B, from left to right: (left) Particles originally picked by Cryolo and used as input for 2D-classification; (middle) the previous set of particles after cleaning by a round of 2D classification (note that discarded particles correspond to those ones belonging to rejected 2D classes, which are marked with a red cross in A); (right) Cryolo original set of particles after application of MicrographCleaner. It can be appreciated that MicrographCleaner removed all particles picked on carbon but 2D-classification did not.

Roughly speaking, our results point out that 2D-clustering is a much more aggressive strategy that removes many more particles that MicrographCleaner (between 20% and 40% compared to 9% to 25%). Obviously, these results should not be surprising as MicrographCleaner was not designed to remove some types of false positive cases (e.g. background) that 2D-classification can.

However, the most interesting conclusions can be drawn when counting the number of particles removed by MicrographCleaner that were not ruled out after 2D-classification (it is acknowledged that particle pruning through 2D classification has a certain subjectivity, difficult to reproduce precisely). Thus, we have measured that between 19% and 29% of the particles discarded by MicrographCleaner survived to the 2D-classification process. Even more interestingly, when a second step 2D-classification is performed, the number of not removed particles, although smaller, is still of consideration (between 10% to 20%, see Supplementary Material S9). These numbers suggest that MicrographCleaner and 2D-classification should better be regarded as complementary options rather than competitors.

## 4. Discussion

Deep learning particle pickers are increasingly gaining popularity. Their ability to avoid contaminated regions and their reported superior accuracies compared to traditional approaches can explain this trend. Yet, traditional particle pickers are still the preferred option in recent publications. Irrespective of the particular method that a researcher considers appropriated for a particular case, we introduce here an approach that is specifically tailored to detect those particles that are located in problematic areas of the micrograph. In other words, rather than concentrating on reporting specimen-like images, we focus on detecting those areas of the micrograph that are likely to contribute with less quality images, so that we can select from any picking method only those particles that are coming from the best areas of the micrograph. Interestingly, we also show how this contextual approach can complement very well other traditional particles selection procedures, such as pruning by 2D classification, in that a quite substantial percentage of images tend to be accepted by 2D classification cleaning that, however, our method detects and discard. Thus, similarly to the general trend in the machine learning field in which top performing solutions are based on ensembles of methods, it is very likely (indeed, it is our vision) that top performing image processing or preprocessing workflows will likely be constructed by combining different approaches, MicrographCleaner included, especially when facing difficult samples.

## 5. Conclusions

MicrographCleaner is an easy-to-install and easy-to-use python package that allows efficient and automatic micrograph segmentation with the aim of preventing particle pickers from selecting inappropriate regions on the micrograph. To that end, MicrographCleaner relays on a U-net-like model that has being trained on about 500 micrographs. When compared to other methodologies, MicrographCleaner has proven more robust, achieving results closer to the human criterion than other methods for both carbon and contaminants detection. In conclusion, we consider that MicrographCleaner is a powerful approach to be applied at the very beginning of cryo-EM workflows, even within on-the-fly/streaming processing pipelines, leading to cleaner sets of input particle and, consequently, to a better processing performance.

## Supporting information

Supplementary sections

## Acknowledgements

The authors would like to acknowledge economical support from: The Spanish Ministry of Economy and Competitiveness through Grants BIO2016-76400-R(AEI/FEDER, UE) and the “Comunidad Autónoma de Madrid” through Grant: S2017/BMD-3817. Ruben Sanchez-Garcia is recipient of a FPU fellowship. The authors acknowledge the support and the use of resources of Instruct-ERIC, a Landmark ESFRI project.

