## Supplementary sections for "MicrographCleaner: a python package for cryo-EM micrograph cleaning using deep learning"

**SUPPLEMENTARY MATERIAL**

### S1. Neural Network Architecture.

A U-net-like (Ronneberger et al., 2015) architecture was employed in this work. Several architectural variations were considered and compared through cross-validation (see S4). The main difference between the classical configuration and ours is that our convolution blocks are indeed residual blocks (Wu et al., 2017) and that we do not perform the crop operation. Other changes are the depth, measured in terms of downsampling operations, which is five instead of four and the size of the input images, which is 256x256 instead 572x572. The number of filters in each convolution operation for each block is 32, 64, 128, 256 and 512 respectively and the filter size is 5x5, except for the first layer, which is 7x7. As activation function, we employed leakyRelu with α=0.05. Batch normalization is also employed. Finally, dropout is added after each residual block of the downsampling part of the network. An scheme of the network architecture can be found in http://campins.cnb.csic.es/micrograph_cleaner/architecture.png and it is summarized in table TS1.1 and TS1.2. The total number of parameters contained in our network is 35,479,712.

Table TS1.1. Neural network architecture of downsampling block number i:

| Layer | Type | Parents | # kernels | Kernel size |
|---|---|---|---|---|
| 1 | Conv2d+BN+LeakyRelu | Previous_block | $2^{i+4}$ | 5x5 (7x7 if i=1) |
| 2 | Conv2d+BN | 1 | $2^{i+4}$ | 5x5 (7x7 if i=1) |
| 3 | Conv2d+BN | Previous_block | $2^{i+4}$ | 1x1 |
| 4 | Add+LeakyRelu+Dropout | 2, 3 | None | None |
| 5 | MaxPooling | 4 | None | 2x2 |

Table TS1.2. Neural network architecture of upsampling block number i:

| Layer | Type | Parents | # kernels | Kernel size |
|---|---|---|---|---|
| 1 | Upsampling2D | Previous_block | None | None |
| 2 | Concatenation | 1, dowmsampling_block_i_l4 | None | None |
| 3 | Conv2d+BN+LeakyRelu | 2 | $2^{N-i+5}$ | 5x5 |
| 4 | Conv2d+BN | 3 | $2^{N-i+5}$ | 5x5 |
| 5 | Conv2d+BN | 2 | $2^{N-i+5}$ | 1x1 |
| 6 | Add+LeakyRelu | 4, 5 | None | None |

### S2. Neural Network Training.

We have employed as loss function the sum of perceptual loss (Johnson et al., 2016) and weighted binary cross-entropy (Falk et al., 2019) at 1:1 proportion. Other alternatives were ruled out after cross-validation (see Section S4). The rationale behind the addition of the perceptual loss term is that for the particular task of contaminants detection, we are not really concerned about the perfect matching of all the pixels, which in large contaminated regions tend to be homogeneous, but about the detection of informative, and thus, more abstract features such as edges that can help the network to better identify the boundaries of the regions.

In order to obtain the perceptual loss estimator, we have trained a classical VGG-Net-16 (Simonyan and Zisserman, 2014) on a grayscale version of the ImageNet dataset (Deng et al., 2009). The VGG-Net-16 trained network is available at http://campins.cnb.csic.es/imagenet_grayscale). Although it is true that the images contained in the ImageNet dataset are of quite different nature compared to micrographs, it is also true that the purpose of the network is to identify carbon and contaminated areas, which are more similar to natural images in terms of signal to noise ratio. Moreover, it is common in the field of deep learning to use networks trained on ImageNet for fine tuning or feature extraction in other domains such as medical imaging (Bar et al., 2015).

Regarding our U-net-like model, Adam optimizer was employed on batches of size 12. L1 and L2 regularization was added to all kernels weights with strength 1e-4. The network was trained until no improvement in validation loss was detected once two learning rate decays on plateau were performed. Initial learning rate was 1e-2.

Severe data augmentation has been performed at a ratio 1:4 using as transformations translations, rotations, contrast and brightness alterations, local blurring and small zooming in/out (with the aim of dealing with slightly different particle size estimations).

### S3. Micrograph preprocessing.

Each micrograph is normalized in intensity by subtracting the median value of its pixels and dividing by the percentile 5-95 range.

$$I_{norm} = \frac{I - median(I)}{percentile_{95}(I) - percentile_{5}(I)}$$

Then, micrographs are downsampled with the aim of normalizing the particle size to 16 pixels. The idea behind this size normalization is that if we are interested in detecting contaminants while preserving particles, having always particles of the same size will simplify the problem. Thus, we apply the following downsampling factor:

$$downFactor = \frac{16}{estimatedParticleWidthInPixels}$$

Thus, the final size of the micrograph is:

$$micrographSize = originalMicrographSize \cdot downFactor$$

When users do not provide an estimation of the particle size, the Scipion “particle boxsize” protocol can be employed to obtain such estimation.

### S4. Architectural decision based on cross-validation

In order to perform cross-validation, the training set was divided into training and validations splits at a proportion 10:1. The following tables summarize the most relevant trials regarding the architecture and other hyperparameters. Metrics included in this table are computed per patch (256x256), contrary to metrics reported in the main text that are computed using the whole micrographs.

Table S4.1. Cross-validation results with respect different numbers of blocks

| Model number | Training mIoU | Validation mIoU | Depth |
|---|---|---|---|
| 1 | 0.6272 | 0.5312 | 4 |
| 2 | 0.6341 | 0.5498 | 5 |
| 3 | 0.6661 | 0.5305 | 6 |
| 4 | 0.6832 | 0.4721 | 7 |

Table S4.2. Cross-validation results with respect different losses

| Model number | Training mIoU | Validation mIoU | Loss |
|---|---|---|---|
| 5 | 0.5991 | 0.4165 | BCE |
| 6 | 0.6523 | 0.5212 | WBCE |
| 7 | 0.6221 | 0.4782 | PL |
| 8 | 0.6341 | 0.5498 | PL+ WBCE |

Notes: BCE: binary cross-entropy; WBCE: weighted binary cross-entropy; PL: perceptual loss.

Table S4.3. Cross-validation results with respect downsampling factor

| Model number | Training mIoU | Validation mIoU | Particle size |
|---|---|---|---|
| 9 | 0.6001 | 0.4799 | 8 |
| 10 | 0.6341 | 0.5498 | 16 |
| 11 | 0.6279 | 0.4812 | 32 |

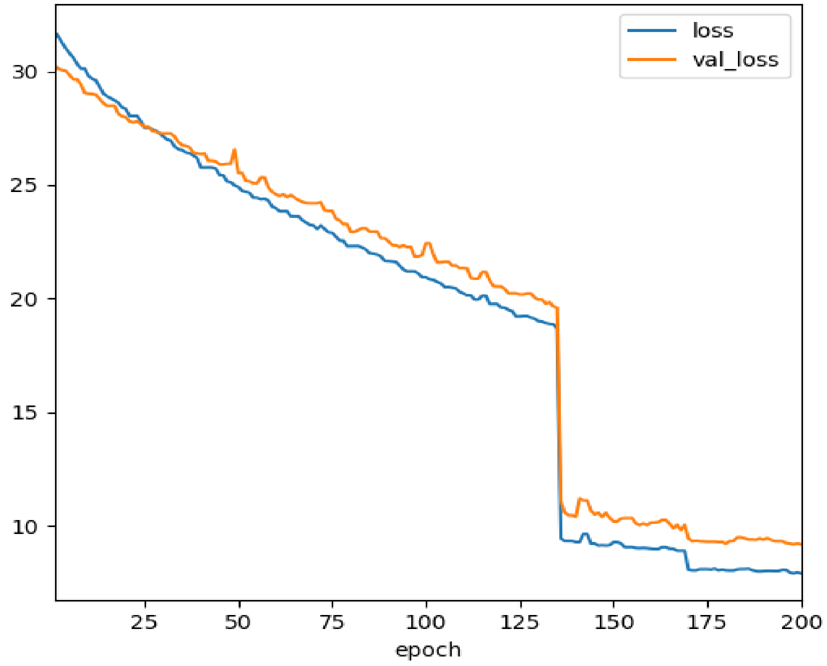


Fig SM1. MicrographCleaner learning curve (loss functions as introduced in S2 above). Loss: training loss, val_loss: validation loss.

### S5. Datasets

#### Evaluation datasets

EMPIAR-10205. Cowpea mosaic virus
EMPIAR-10217. Bovine liver glutamate dehydrogenase
In-home dataset1. Phage T7 tails
In-home dataset2. Phage T7 ejection machinery

#### Training datasets

EMPIAR-10005. TRPV1
EMPIAR-10028. Ribosome
EMPIAR-10033. Picornavirus
EMPIAR-10049. RAG1-RAG2 Complex
EMPIAR-10061. β-galactosidase
EMPIAR-10075. Phage MS2
EMPIAR-10077. Elongation factor SelB
EMPIAR-10081. HCN1 ion channel
EMPIAR-10090. AAA-ATPase in 26S proteasome
EMPIAR-10093. Ion channel in nano disc
EMPIAR-10097. Influenza Hemagglutinin Trimer
EMPIAR-10099. Hrd1 and Hrd3 complex
EMPIAR-10168. RNA Polymerase III pre-initializ
EMPIAR-10175. Hemagglutinin
EMPIAR-10190. RNA Polymerase III transcribing
EMPIAR-10203. Nodavirus

### S6. Carbon detection capability comparison

In order to compare MicrographCleaner carbon detection capability with em_hole_finder, EMHP and the Warp particle picker (WPP) BoxNet2Mask_20180918 model, we have executed the four software packages on the same testing dataset consisting of a subset of 60 micrographs, all them containing carbon. Contrary to MicrographCleaner and WPP, EMHP and em_hole_finder require from the user to set certain parameters. For this comparison, we have set those parameters to default values. While custom tuning of them should improve results, we have adopted this approach because of two reasons. First, we have tried different parameters settings and we have not observed too important performance differences on average, since some settings caused some inputs to improve at the cost of worsening others. Secondly, we believe that solutions that require considerable human intervention are in danger of extinction as the cryo-EM field is moving towards streaming and automatic processing and thus, default parameters should perform decently for most cases. Other methods used in computer vision (k-mean clustering and SLIC), where initially considered for comparison purposes, but were discarded for the reasons presents above, particularly, because of the difficulty of setting parameters and thresholds that could be valid for all the elements of the dataset.
The quality of the predicted masks has been assessed by comparing them to ground truth masks manually compiled. In the case of MicrographCleaner and WPP, as they also find contaminated regions, we have only considered the predicted patches that overlapped with the ground truth carbon masks (if they exist), ignoring predictions for contaminated regions. As shown in Main Text Table 1, MicrographCleaner is able to produce predictions much more similar to the ground truth masks than the other methods, with a mean Intersection over Union (mIoU) value close to 0.8. The second-best performing algorithm is WPP, which does a much better job than EMHP and em_hole_finder, although still far from MicrographCleaner performance. The small mean mIoU obtained by EMHP and, especially, by em_hole_finder, can be explained by the lower quality masks they produced when compared with MicrographCleaner and, mainly, by the number of total failure cases that both methods suffers, that is, cases in which the overlapping between the ground truth and the predicted mask is 0 (see Figure SM1 E). Figure SM1 shows and example of success in which all the four algorithms are able to detect, to an acceptable extend, the

carbon present in the micrographs (superior row images) and an example of failure, in which only MicrographCleaner and WPP have been able to detect the carbon.

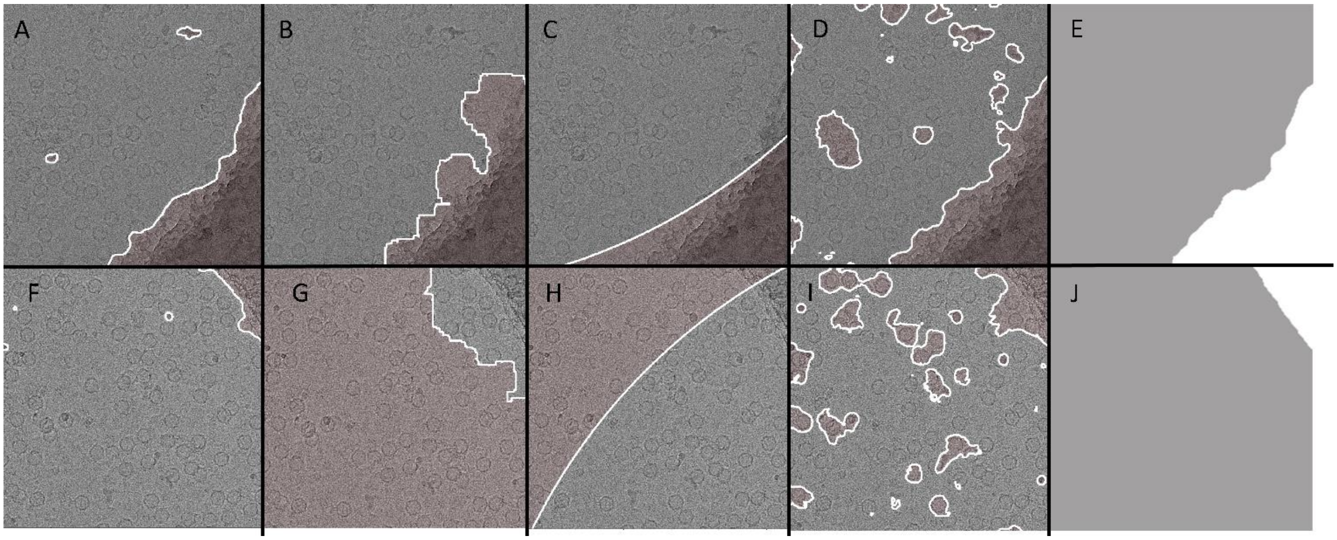


Figure SM2. Images A-D display the predicted masks (red shadowed areas) obtained with MicrographCleaner, em_hole_finder, EMHP and WPP, respectively, on the same micrograph. Image E shows the manually curated mask used to evaluate carbon detection. In this case, all algorithms have been able to obtain acceptable solutions. Images F-I display the predicted masks (red shadowed areas) obtained with MicrographCleaner, em_hole_finder, EMHP and WPP respectively, on another micrograph. Similarly, image J displays the manually curated mask for evaluation. Figure G represents an example of total failure in which not even a single carbon pixel was detected.

### S7. Undesirable regions detection

Additionally, we have compared MicrographCleaner and WPP segmentation capability, measuring the mIoU of the predictions of both algorithms for all the micrographs contained in the testing set. Thus, MicrographCleaner has being able to obtain better predictions than WPP for 77.66% of the micrographs used for evaluation and, over all, it has achieved a mIoU of 0.544 as compared to the 0.331 value measured for WPP. Figure SM2 shows the predictions obtained for four different micrographs using MicrographCleaner (bottom) and WPP (top). As it can be appreciated in these results, MicrographCleaner predictions are, generally speaking, better fitted to the actual contaminants present in the micrographs. Nevertheless, WPP is able to perform better than MicrographCleaner for 22.33% of the evaluated cases (as illustrated in SM2 D) and thus, both approaches could be jointly considered to obtain better results.

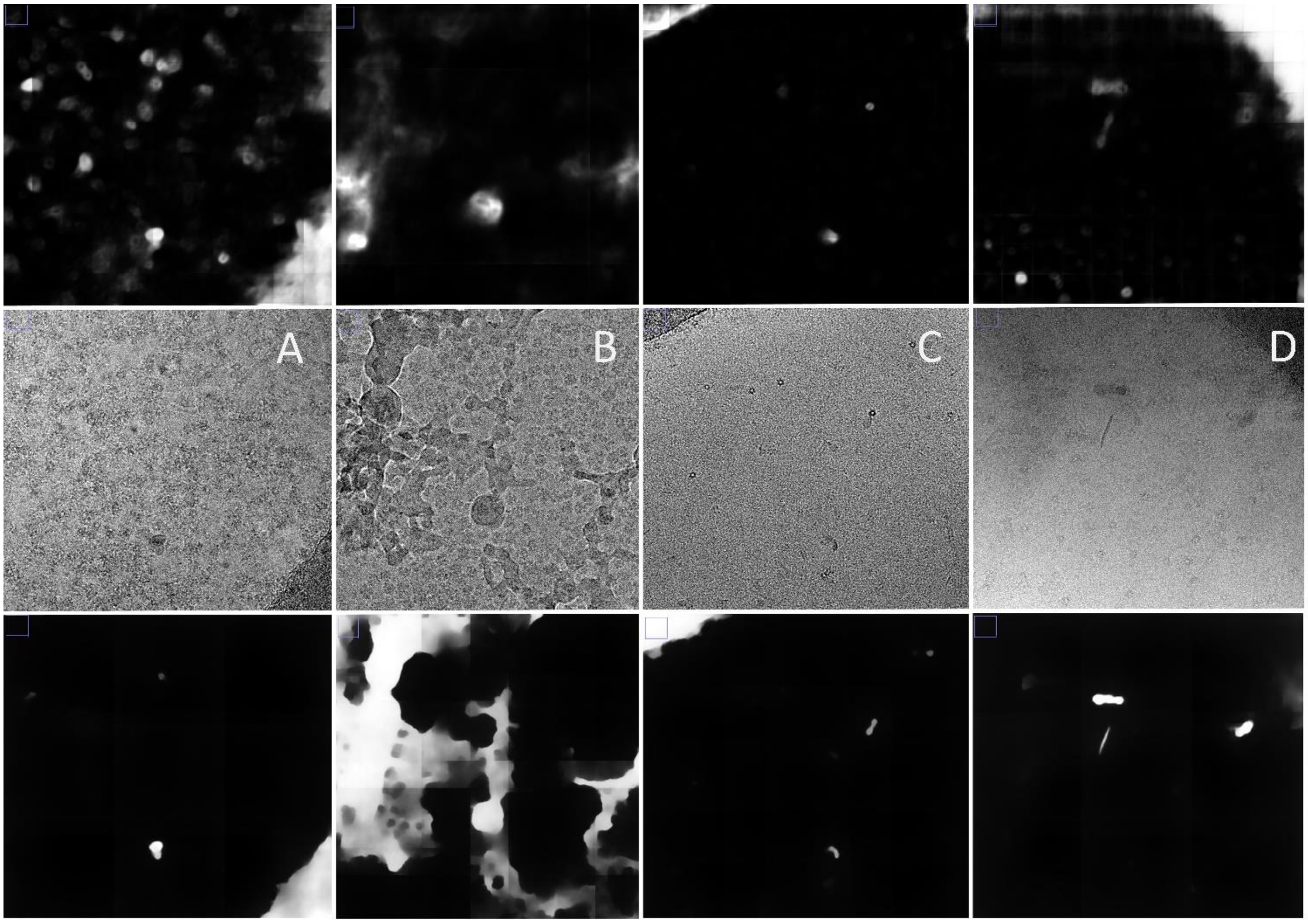


Figure SM3. Four different examples (A-D) computed using MicrographCleaner (bottom row) and the Warp particle picker (top row). Predictions for the examples A and B are clearly favorable to MicrographCleaner, whereas example C predictions are comparable for both algorithms. Example D compares favorably to Warp particle picking algorithm.

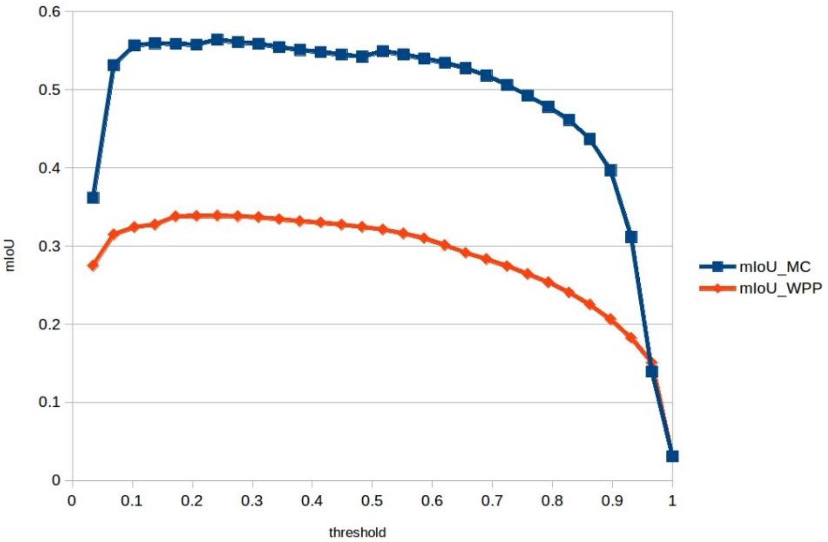


Figure SM4. Performance of MicrographCleaner and the Warp particle picker over the micrographs of the testing set for different threshold options. As it can be appreciated, both approaches are quite robust to threshold selection.

### S8. Use cases

The use cases included in this publication aim to illustrate how MicrographCleaner can boost particle picking no matter the type of used algorithm. The fundament behind this claim is the “No Free Launch” theorem (Wolpert, 1996; Wolpert and Macready, 1997) which implies that there is no a single algorithm better than all others for every [optimization] problem. Although, generally speaking, deep-learning particle pickers tend to outperform classical ones, these examples show some cases in which their performance is not extraordinary and it may be comparable to classical particle pickers. We also show that using an orthogonal method to the pickers (a segmentation method instead of a detection/classification one) we can improve their results. This strategy of employing a blend of different algorithms to improve performance is widely used in the field of machine learning, in which real life solutions tend to be based on boosting and stacking of models and it may be a wise strategy for the cryo-EM specimens that are hard to pick.

In both examples, we have employed the Cryolo particle picker (version 1.5.4) using the general model provided by the authors (version gmodel_phosnet_201910) and also, we have trained a custom model using 10 micrographs manually picked. The Topaz particle picker was also trained using the same micrographs. On the contrary, the Relion autopicker was executed using default parameters. Topaz and Cryolo solutions were manually examined in order to select an adequate global threshold. An alternative stricter threshold was also manually selected with the aim of removing most of the detected particles in the carbon area/edges as well as the contaminants picked as particles.

*S.8.1. EMPIAR-10156*

None of the particle pickers studied was able to perfectly avoid the carbon regions/edges at reasonable thresholds. Main Text Figure 2 is one of the many examples in which a non-negligible number of false positive particles are selected independently of the algorithm and the threshold. Generally speaking, all the particle pickers are able to select most of the true positive particles but the carbon areas are not being avoided by the Relion particle picker and Cryolo is also missing some of them. On the contrary, the Topaz algorithm is able to better avoid the carbon regions, although it is strongly attracted by the carbon edges, which are equally if not more dangerous.

When comparing the total number of picked particles (Table S.8.1), it can be appreciated that the Cryolo particle picker solution is the one which is less modified by MicrographCleaner and thus it seems to be the cleaner set of particles. On the contrary, the Relion autopicker is the one that is more affected by MicrographCleaner, which supports the idea that traditional particle pickers are more affected by false positives than deep-learning-based solutions. However, after MicrographCleaning usage, all the different approaches contain a similar number of particles, which suggests that the quality of the datasets might be comparable. This should not be so surprising as the picking of the particles in this example is not complicate leaving aside the problem of the carbon areas/edges.

Table S.8.1. Number of picked particles using different algorithms and thresholds.

| Algorithm | Threshold | #particles |
|---|---|---|
| Cryolo general model | 0.1 | 13916 |
| Cryolo general model + MicrographCleaner | 0.1 | 12620 |
| Cryolo general model | 0.3 | 9801 |
| Cryolo trained | 0.4 | 14262 |
| Cryolo trained+ MicrographCleaner | 0.4 | 13211 |
| Cryolo trained | 0.5 | 10013 |
| Topaz | 2 | 17981 |

| | | |
|---|---|---|
| Topaz + MicrographCleaner | 2 | 15321 |
| Topaz | 4 | 13592 |
| Relion auto | Default | 19616 |
| Relion auto + MicrographCleaner | Default | 14852 |

*S.8.2. EMPIAR*-10265

In this example, and contrary to the previous one, the Cryolo solutions, specially the one obtained from the manually trained model is substantially better that the others. Yet, it is still not perfect as illustrated by the small contaminants displayed in Main Text Figure 3 and 4 and by the fact that MicrographCleaner is still able to remove more than 2000 false positive particles from them. Hence, and likewise the previous case, if most of the contaminants that can visually be identified are discarded using a higher threshold, there is still an important number of true positive particles that are also removed, as it can be derived from the numbers exposed in Table S.8.2. However, these effects are less pronounced that for the other methods.

Regarding the Topaz and the Relion solutions, it is important to notice that in this case, the number of discarded particles, 6% and 15% respectively, is of importance. It might seem surprising that a relatively small set of discarded images make a difference in the quality of downstream analysis of cryEM particles, but we should note that there is a very profound statistical difference between adding or removing "randomly" a set of particles, and adding or removing a totally biased set of particles (those in bad micrograph areas). This is an area in which we anticipate substantial developments in the cryoEM area be coming: Being aware of bias in the calculation of cryoEM maps, and introducing the appropriate means of correction, as we are doing here wih MicrographCleaner.

Table S.8.2. Number of picked particles using different algorithms and thresholds.

| Algorithm | Threshold | #particles |
|---|---|---|
| Cryolo general model | 0.1 | 168820 |
| Cryolo general model + MicrographCleaner | 0.1 | 165070 |
| Cryolo general model | 0.3 | 97085 |
| Cryolo trained | 0.5 | 203131 |
| Cryolo trained+ MicrographCleaner | 0.5 | 198383 |
| Cryolo trained | 0.7 | 104121 |
| Topaz | 3 | 140204 |
| Topaz + MicrographCleaner | 3 | 131572 |
| Topaz | 5 | 63843 |
| Relion auto | Default | 177029 |
| Relion auto + MicrographCleaner | Default | 149720 |

### S9. MicrographCleaner complements 2D-classification

In order to study the effectiveness of MicrographCleaner when 2D-classification is considered, we have run Relion 2D-classification (Kimanius et al., 2016; Scheres, 2012) algorithm on the sets of particles that were presented in Main Text section 3.3.1 and Supplementary Material S8.1. Then, we have ruled out all the particles that belonged to "bad classes" and we have counted the number of particles that remained. Additionally, we have computed the intersection between the sets of particles that remained after 2D-classification processing and the particles removed by MicrographCleaner. As the number of particles discarded by MicrographCleaner that are not removed by 2D-classification was large (19% to 29%), we performed a second round of 2D-classification in order to clean further the sets of particles. Yet, the

number of particles ruled out by MicrographCleaner that survived after two rounds of classification, as illustrated in table S.9, is important, between 10% to 19%.
Along the next pages, we present the same example than Main Text Figure 5 when the remaining particle pickers are considered instead. In all them, the situation is similar to Main Text Figure 5, as MicrographCleaner is able to remove the particles that lay on the carbon/edges while the 2D-classification algorithm is not able to fully remove them.

Table S.9. Number of remaining particles after 2D classification and MicrocraphCleaner in *EMPIAR-10156*

| Algorithm | Threshold | initial | 2D-classes$_1$ | micClean | 2D-classes$_1$ & micClean removed | 2D-classes$_2$ | 2D-classes$_1$ & micClean removed |
|---|---|---|---|---|---|---|---|
| Cryolo general | 0.1 | 13916 | 11045 | 12620 | 377 | 6776 | 205 |
| Cryolo trained | 0.4 | 14626 | 8873 | 13211 | 329 | 7021 | 273 |
| Topaz | 2 | 17981 | 12046 | 15321 | 498 | 6876 | 253 |
| Relion auto | NA | 19616 | 12554 | 14852 | 1055 | 7941 | 608 |

NOTE: initial: Number of particles that were selected by the particle picker; 2D-classes$_1$: Number of particles that survived first step 2D-classifaction; micClean: Number of particles that survived to MicrographCleaner execution; 2D-classes$_1$ & micClean removed: number of particles that survived to the first step of 2D-classification but are false positives according to MicrographCleaner

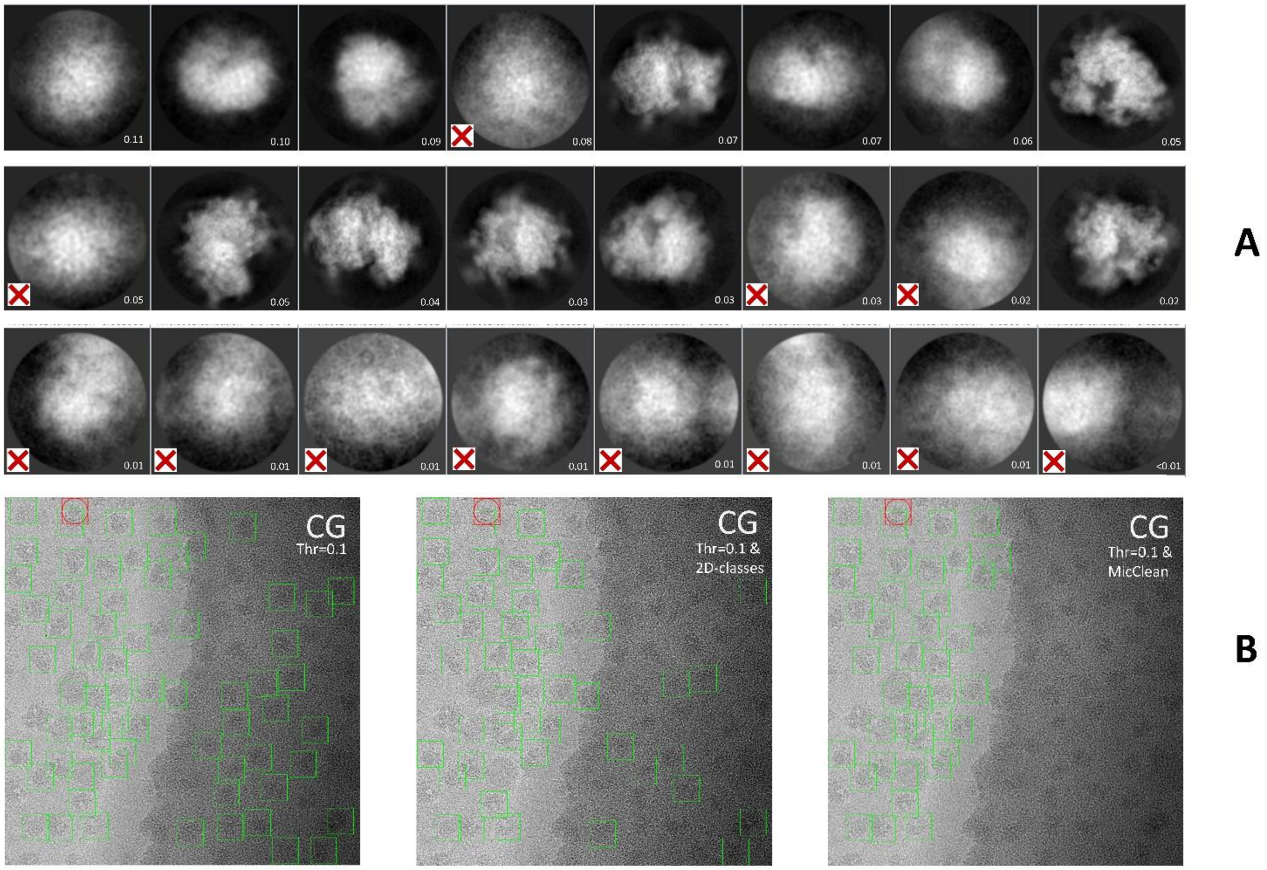


Figure SM5. MicrographCleaner complements 2D-classification. A: Gallery of 2D averages obtained from the set of particles collected by Cryolo general model on EMPIAR-10265 dataset. B, from left to right: (left) Particles originally picked by Cryolo and used as input for 2D-classification; (middle) the previous set of particles after cleaning by a round of 2D classification (note that discarded particles correspond to those ones belonging to rejected 2D classes, which are marked with a red cross in A); (right) Cryolo original set of particles after application of MicrographCleaner. It can be appreciated that MicrographCleaner removed all particles picked on carbon but 2D-classification did not.

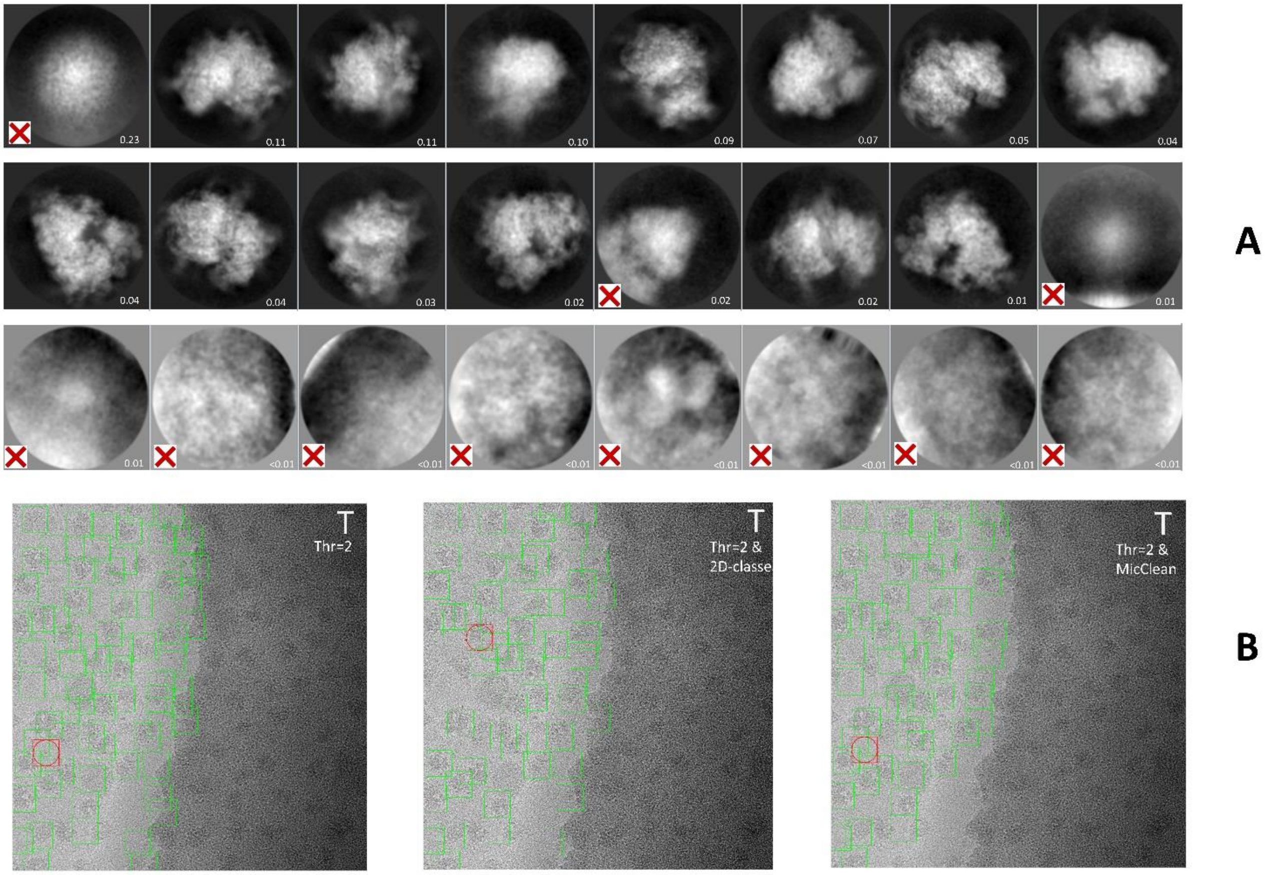


Figure SM6. MicrographCleaner complements 2D-classification. A: Gallery of 2D averages obtained from the set of particles collected by Topaz on EMPIAR-10265 dataset. B, from left to right: (left) Particles originally picked by Topaz and used as input for 2D-classification; (middle) the previous set of particles after cleaning by a round of 2D classification (note that discarded particles correspond to those ones belonging to rejected 2D classes, which are marked with a red cross in A); (right) Topaz original set of particles after application of MicrographCleaner. It can be appreciated that MicrographCleaner removed all particles picked on carbon but 2D-classification did not.

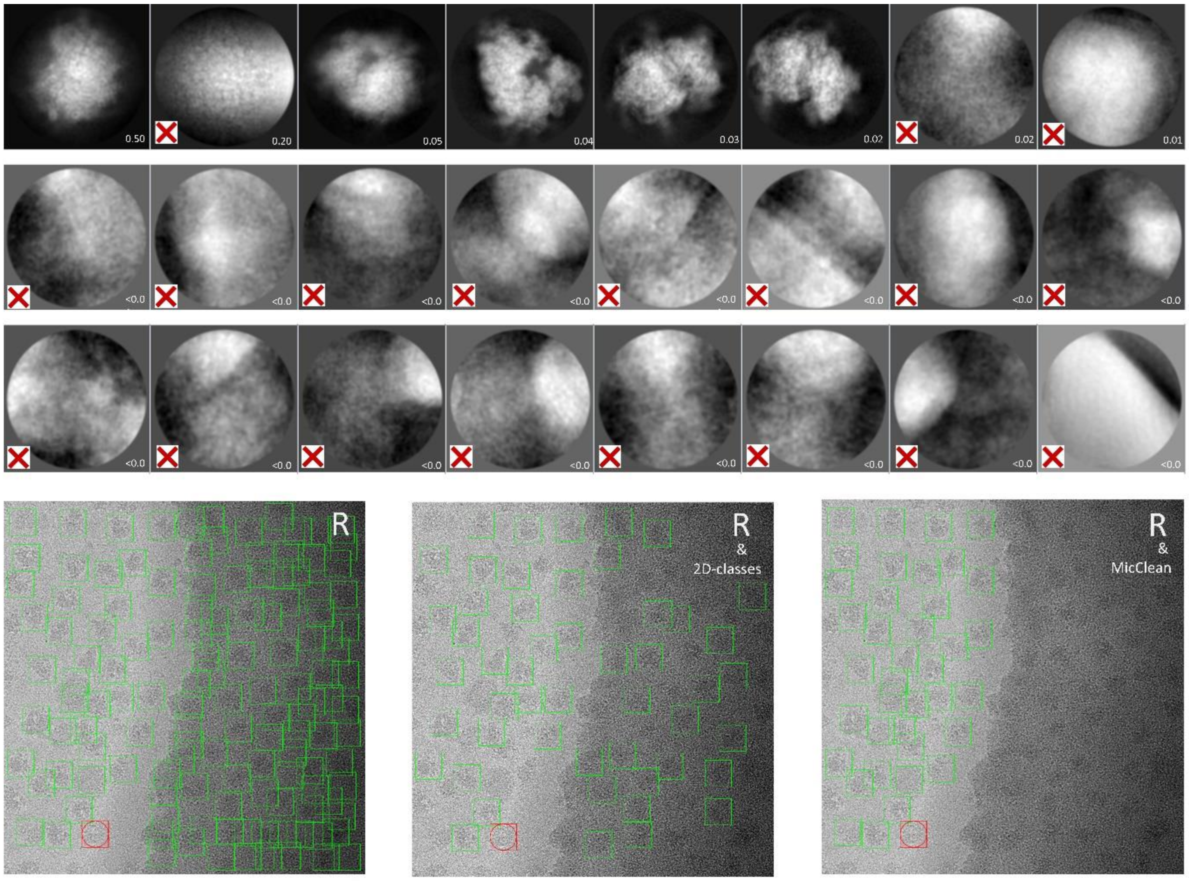


Figure SM7. MicrographCleaner complements 2D-classification. A: Gallery of 2D averages obtained from the set of particles collected by Relion autopicker on EMPIAR-10265 dataset. B, from left to right: (left) Particles originally picked by Relion autopicker and used as input for 2D-classification; (middle) the previous set of particles after cleaning by a round of 2D classification (note that discarded particles correspond to those ones belonging to rejected 2D classes, which are marked with a red cross in A); (right) Relion autopicker original set of particles after application of MicrographCleaner. It can be appreciated that MicrographCleaner removed all particles picked on carbon but 2D-classification did not.

### S10. Usage guide

A complete installation and command line execution guide can be found in https://github.com/rsanchezgarc/micrograph_cleaner_em/tree/master.

In order to compute masks from micrographs, just two commands are needed to be employed once the package is installed.

First, with the aim of downloading an updated version of the deep learning model, the following command should be executed:

```
cleanMics --download
```

Then, in order to compute masks for a given set of micrographs, the following command should be executed:

```
cleanMics -b $BOX_SIXE -i /path/to/micrographs/ --predictedMaskDir path/to/store/masks
```

Additionally, MicrographCleaner can also be executed within the cryo-EM framework Scipion (de la Rosa-Trevín et al., 2016) through the protocol deepMicrographScreen. An illustration of the form is depicted in Figure SM8.

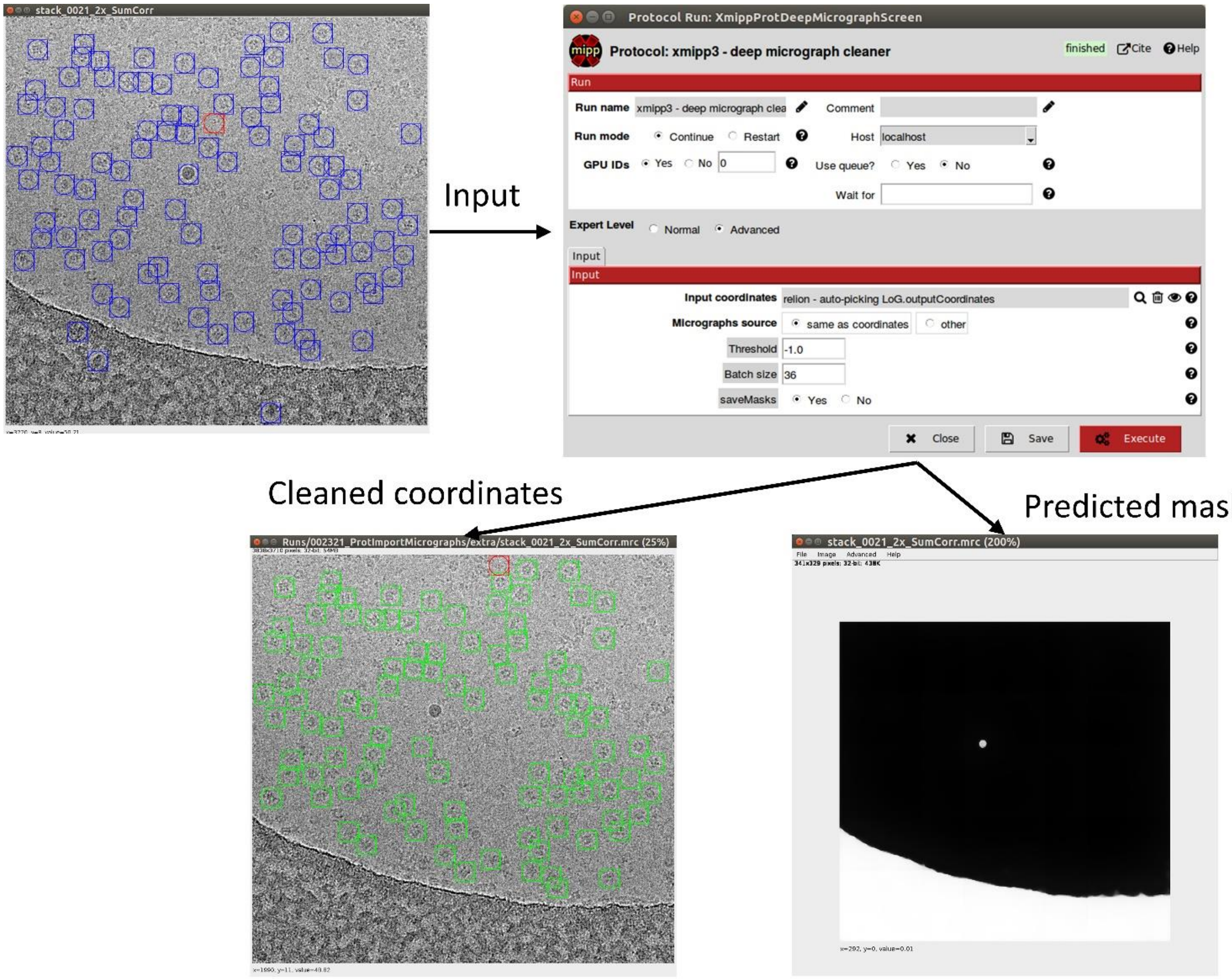


Figure SM8. MicrographCleaner GUI illustration within Scipion.

As it can be appreciated in SM8, the only two parameters required for the execution of MicrographCleaner scipion protocol are the batch size, that only has impact over the GPU computation efficiency, and the threshold, that can be manually set or left with the default value.

Additionally, MicrographCleaner can be programmatically executed, and thus, integrated with other tools, using a simple API documented in https://github.com/rsanchezgarc/micrograph_cleaner_em/tree/master.
